# Spreading depolarization and repolarization during cardiac arrest as an ultra-early marker of neurological recovery in a preclinical model

**DOI:** 10.1101/786210

**Authors:** Robert H. Wilson, Christian Crouzet, Donald E. Lee, Dishant P. Donga, Ayushi H. Patel, Afsheen Bazrafkan, Niki Maki, Masih A. Rafi, Maziar Moslehyazdi, Justin H. Pham, Mohammad Torabzadeh, Brooke E. Hjelm, Bruce J. Tromberg, Oswald Steward, Beth A. Lopour, Bernard Choi, Yama Akbari

## Abstract

Spreading depolarization (SD) accompanies numerous neurological conditions, including migraine, stroke, and traumatic brain injury. There is significant interest in understanding the relationship between SD and neuronal injury. However, characteristics underlying SD and repolarization (RP) induced by global cerebral ischemia (e.g., cardiac arrest (CA)) and reperfusion are not well understood. Quantifying features of SD and RP during CA and cardiopulmonary resuscitation (CPR) may provide important metrics for diagnosis and prognosis of neurological injury from hypoxia-ischemia. We characterized SD and RP in a rodent model of asphyxial CA+CPR using a multimodal platform including electrocorticography (ECoG) and optical imaging. We detected SD and RP by (1) alternating current (AC), (2) direct current (DC), and (3) optical imaging of spreading ischemia, spreading edema, and vasoconstriction. Earlier SD (r=−0.80; p<0.001) and earlier RP (r=−0.71, p<0.001) were associated with better neurological recovery after 24hrs. SD+RP onset times predicted good vs poor neurological recovery with 82% sensitivity and 91% specificity. To our knowledge, this is the first preclinical study to link SD and RP characteristics with neurological recovery post-CA. These data suggest that SD and RP may be ultra-early, real-time prognostic markers of post-CA outcome, meriting further investigation into translational implications during global cerebral ischemia.

## Introduction

Spreading depolarization (SD) is a wave of neuronal depolarization observed in a variety of conditions, including migraine, traumatic brain injury, and stroke.^1,2^ Leão first described the closely related spreading depression phenomena in 1944^3^, and subsequent studies characterized hallmarks of SD, such as shifts in direct current (DC) potential and spreading oligemia.^4–11^ However, SD’s role in neuronal injury and its clinical relevance were not widely recognized until recently.^12^

While SD is commonly associated with migraine and acute to subacute focal brain injuries, SD and its features on electrocorticography (ECoG) are not widely known for global cerebral ischemia. The DC potential shift from SD appears as an ultra-slow wave in low-pass (LP) ECoG in rats.^13,14^ Previous studies characterized this ECoG wave as a “wave of death”.^14,15^ However, this “wave of death” has been shown to merely mark the onset of cytotoxic events (e.g., glutamate release, Ca^2++^ influx, cytotoxic edema) from which the brain can recover with timely reperfusion.^16–18^

Recently, SD’s role in neuronal injury has become a topic of increasing interest in critical care management after acute brain injury.^19^ Recently, SD has been characterized with ECoG in a variety of brain-injured patients at multiple neurosurgical centers.^19–23^ Additionally, SD was recently characterized in cardiac arrest (CA) patients for the first time with multimodal monitoring.^16^

However, the role of SD in neurological outcome has not been investigated in CA and global ischemia. Anoxia-induced SD is often called anoxic depolarization or, if occurring during cardiac arrest or brain death, terminal spreading depolarization, both of which have significant overlap with the various SD phenomena, including spreading and non-spreading depolarization and depression.^1,16,24,25^

For simplicity, in this paper, we will refer to this depolarization during CA as SD. Quantifying SD during CA and resuscitation may provide an important tool for diagnosis, prognosis, and possible therapeutic interventions during neurological injury during hypoxic-ischemic events. Currently, post-CA prognostication within the first 24-72 hours is based on serial neurological exams, including electroencephalography (EEG), imaging (CT and/or MRI), and blood biomarkers such as neuron-specific enolase (NSE).^26^ However, many of the processes responsible for ischemic damage and reperfusion injury in the brain are well-underway by the time that these exams are completed.^27^ Therefore, there remains a critical need for rapid prognostic tools to guide novel therapeutic strategies at ultra-early time points during CA and immediately following resuscitation.

Traditionally, studies have primarily measured SD using DC electrodes or micropipettes.^28^ More recently, several groups have used optical imaging to measure and characterize SD in animal models of cerebral ischemia.^29–32^ Optical methods can also quantify ischemia-induced mismatches between cerebral blood flow (CBF) and brain metabolism that are indicative of impaired autoregulation.^33–39^ Additionally, via measurement of tissue scattering, optical techniques can also interrogate neuronal damage caused by cytotoxic edema and dendritic beading.^24,40^

In this study, we used a preclinical model of asphyxial CA (ACA) and cardiopulmonary resuscitation (CPR),^41,42^ along with an ECoG and optical imaging platform^43,44^, to characterize SD during ACA and repolarization (RP) post-CPR. We show that earlier SD and earlier RP are both associated with better neurological outcome. Additionally, we characterize cerebral perfusion-metabolism dynamics during ACA and post-resuscitation that may be critical for understanding SD, RP, and the recovery of the brain from hypoxia-ischemia. Previous studies have employed hypo- and hyperglycemia interventions to assess the effect on SD susceptibility in global ischemia^45^ while others have tested hyperglycemia’s effects on SD and RP susceptibility in focal and forebrain ischemia.^46,47^ However, to our knowledge, this report is the first observational study to associate SD and RP susceptibility with neurological outcome after global ischemia, and also the first to characterize RP features after reperfusion or resuscitation on alternating current (AC)-ECoG.

## Materials and Methods

### Preclinical Model

The animal model of ACA and resuscitation employed in this study has been described previously^41–44,48^ and is summarized in Fig. 1. Briefly, male Wistar rats (Charles River, Canada facility) were handled daily for at least 1 week prior to any experimentation. The night prior to ACA experiments, rats were calorically restricted with 3 food pellets, as standard with our CA model. The next morning, rats were intubated and received femoral artery and vein cannulation to enable arterial blood pressure (BP) monitoring and drug delivery, respectively. During the ACA experiment (Fig. 1A), isoflurane was washed out over a 3 min period and a paralytic agent (vecuronium) was given. Following the washout period, CA was induced by turning off the ventilator for 5, 7, or 8 min, depending on experimental cohort. Subsequently, CPR was initiated by turning the ventilator back on, giving epinephrine and sodium bicarbonate, and performing manual chest compressions. Chest compressions were continued until return of spontaneous circulation (ROSC), after which the rats were monitored continuously for an additional 2-4 hours.

**Figure 1.**
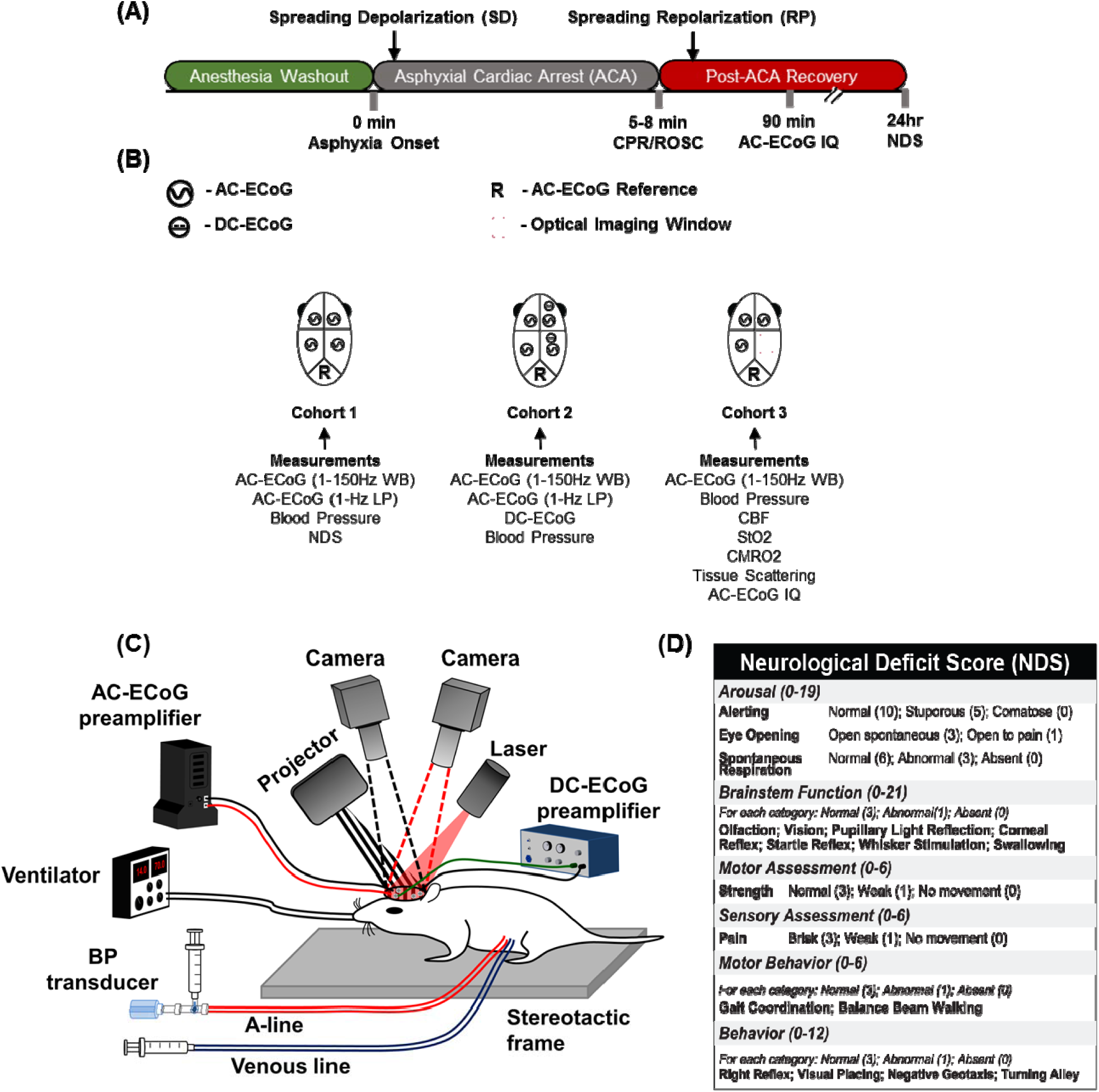
Asphyxial cardiac arrest (ACA) and CPR rat model. (A) Timeline of ACA and CPR experiment. Isoflurane anesthesia is washed out over 3 min, during which a neuromuscular blocker (vecuronium) is administered. ACA is induced by turning off the ventilator for a 5-8 min asphyxial period, depending on experimental cohort. CPR is administered until return of spontaneous circulation (ROSC). Neurological outcome is determined by 24-hr post-ROSC neurological deficit score (NDS) testing or 90-min post-ROSC alternating current electrocorticography (AC-ECoG) information quantity (IQ). (B) Lists of measurement modalities, along with schematics of electrode placement for AC-ECoG and direct current (DC) ECoG, for all three cohorts. (C) Animal model of ACA and CPR for Cohort 3, measured with multimodal monitoring platform. This platform includes arterial-line blood pressure (BP), AC-ECoG, laser speckle imaging (LSI) for measuring cerebral blood flow (CBF), and spatial frequency domain imaging (SFDI) for assessing tissue oxygenation (StO2) and scattering. The combination of LSI and SFDI enables measurement of cerebral metabolic rate of oxygen (CMRO2). (D) Rubric for NDS used to assess neurological recovery 24 hrs post-ROSC. 70 = best outcome, 0 = worst outcome.

In this study, three separate cohorts of rats, each with different multimodal monitoring techniques, were used (Fig. 1B). In Cohort 1, surgery to implant screw electrodes for AC-ECoG recordings was done 1 week prior to the CA experiment. All rats underwent an 8 min ACA and received neurological testing (Neurological Deficit Score; NDS) at 24 hrs post-ROSC (Fig. 1D). Rats in Cohort 2 also underwent 8 min ACA, and had two additional ECoG silver chloride (Ag/AgCl) electrodes implanted to record DC potential.^13^ The Ag/AgCl electrodes were made by chloridizing silver wires (Stoelting 50880) in household bleach. The Ag/AgCl electrodes were implanted 1 mm into the cortex in the right hemisphere, within 3 mm of the right occipital AC-ECoG screw electrodes.

Cohort 3 was studied with optical imaging (Fig. 1C) after performing a partial craniectomy over the right parietal lobe on the day of the experiment. Rats in this cohort were monitored continuously with 3 AC-ECoG electrodes, but did not have DC-ECoG monitoring. In addition, these animals were split into a “mild CA” (5 min asphyxia) and a “moderate CA” (7 min asphyxia) group to mimic the variability of CA durations encountered in an intensive care setting. Rats in this cohort were monitored for 2 hrs post-ROSC and then sacrificed, as the presence of the craniectomy prevented them from being survived. 24-hr NDS was unable to be performed on Cohort 3, so 90 min post-ROSC ECoG information quantity (IQ) was used to measure short-term neurological outcome following resuscitation.^49^

### Data Acquisition and Post-Processing

Post-processing of arterial BP, AC- and DC-ECoG, and optical imaging data was performed using MATLAB (The MathWorks, Inc., Natick, MA).

#### Arterial BP and Arterial Blood Gas (ABG)

For all cohorts, BP was measured continuously at 191 Hz from an arterial line. Mean arterial pressure (MAP) was determined at 1-Hz. ABGs were acquired prior to start of experiment with i-STAT 1 Analyzer (Abaxis, Union City, CA).

#### AC/DC-ECoG

For all cohorts, AC-ECoG data was acquired at 1526 Hz with the RZ5D BioAmp Processor (Tucker-Davis Technologies, Alachua, FL) and PZ2 preamplifier with inbuilt 0.35 Hz – 7.5 kHz band-pass filter. In Cohort 2, in parallel with AC-ECoG data, DC-ECoG data was acquired at 305 Hz with Duo 773 and Model 750 electrometers (World Precision Instruments, Saratosa, FL).

For wide-band (WB) ECoG, AC-ECoG data was 60-Hz notch filtered and 1-150Hz band-pass filtered. To visualize SD on AC-ECoG, a 1-Hz LP filter was applied. A 1-Hz LP filter was also applied to DC-ECoG data to remove noise.

For Cohort 3, the median ECoG IQ from 90 to 100 min post-ROSC was calculated from WB ECoG, as previously described.^49^ The baseline microstates were determined from the final minute of anesthesia washout prior to start of asphyxia. AC-ECoG IQ was also normalized to this baseline period.

#### Optical Imaging

Rats in Cohort 3 underwent continuous laser speckle imaging (LSI) and multispectral spatial frequency domain imaging (SFDI) throughout the experiment. Both LSI and SFDI utilize diffuse near-infrared light to interrogate subsurface tissue structure and function. LSI is an established technique to measure blood flow in preclinical models and humans,^50–52^ including CA studies.^44,53,54^ SFDI is an established technique ^55,56^ that enables separate quantification of the diffuse optical absorption and scattering coefficients of tissue. The absorption coefficient, when measured at multiple near-infrared wavelengths, provides information regarding concentrations of oxygenated and deoxygenated hemoglobin (ctHbO2, ctHb, respectively) in tissue.^43,56,57^ The scattering coefficient provides information about structure, distribution, and concentration of tissue components, such as cell membranes, nuclei, and mitochondria.^40^ Increased tissue scattering is a known indicator of cytotoxic edema that accompanies SD.^24^ Combining cerebral blood flow (CBF) data from LSI with cerebral oxygenation data from SFDI enables calculation of cerebral metabolic rate of oxygen consumption (CMRO2).

### Data Analysis

#### Exclusion Criteria

For Cohort 1, three rats were excluded from analysis due to poor quality AC-ECoG recordings. SD was observed in n=27 rats. RP onset was unable to be determined in 5 rats, so n=22 for analysis involving RP onset. No rats were excluded from Cohort 2 (n=5). For Cohort 3, one rat was excluded due to prolonged CPR and blood loss complications. Thus, for LSI data n=10 (5-min ACA = 5, 7-min ACA = 5). One additional rat was excluded from SFDI analysis due to an instrument malfunction. Thus, for SFDI data n=9 (5-min ACA = 4; 7-min ACA = 5).

#### LSI Data: Speed and Spatial Propagation of Spreading Ischemia

To characterize the CBF waves, videos of CBF images were created. Each frame of the video consisted of CBF images averaged over one-second intervals. The videos were used to characterize the periods of spreading CBF waves. The spatial onset and completion locations of each wave and their corresponding times were extracted after visual inspection of each video. To calculate the speed of the wave, the distance between the onset and completion locations and the duration of the waves were used. To quantify the total amount of brain perfusion prior to the onset of spreading ischemia, we integrated over time the relative CBF (rCBF) time-course signal from the onset of asphyxia to the onset of spreading ischemia. rCBF was normalized to the mean CBF calculated over the one-minute interval immediately prior to the onset of asphyxia.

#### LSI Data: Vessel Diameter

Custom-written MATLAB code was used to compute vessel diameter using one second-averaged CBF images. First, the centerline for a given vessel of interest was defined, followed by the bounds for a line that was perpendicular to the centerline. For each time point, the CBF values from the perpendicular line were extracted, and a Gaussian was fitted through the CBF values. If the R-squared of the Gaussian fit was greater than 0.9, the full-width half-maximum (FWHM) was computed from the Gaussian fit. If the value was less than 0.9, the diameter was assigned a not-a-number value. This process was repeated on 5 to 10 vessels per experiment.

#### SFDI Data: Tissue scattering and oxygenation

Custom-written MATLAB code^43^ based on previous SFDI work developed at Beckman Laser Institute^56^ was used to fit a photon propagation model (Monte Carlo simulation) to the data to extract maps of the tissue absorption and scattering coefficients at the three measured wavelengths (655 nm, 730 nm, 850 nm). The absorption coefficient was then fit to a linear combination of oxygenated and deoxygenated hemoglobin absorption to extract the concentrations of oxy- and deoxy-hemoglobin (ctHbO2, ctHb) and the tissue oxygenation (StO2). To generate ctHbO2, ctHb, StO2, and tissue scattering time-courses, a region of interest (ROI) over the parenchyma (i.e., not atop a major vessel) was selected for each individual rat. Each rat’s ROI was kept constant for every time point in an experiment.

#### LSI + SFDI Data: CMRO2

To generate CMRO2 time-courses, LSI data was combined with SFDI data, following the method of Boas and Dunn.^58,59^ For LSI data, an ROI covering nearly the entire craniectomy was used. For SFDI data, an ROI atop a vein was used when calculating ctHb and ctHbO2. Note that the CMRO2 calculation used a different ROI for the SFDI data than the initial scattering and absorption calculation. This is because the ctHb and ctHbO2 in a venous ROI were considered a better indicator of the oxygen consumed by the brain, an important distinction for the CMRO2 formula.

#### Multimodal Parameter Set

For each cohort, a multimodal set of parameters was extracted from the data pre- and post-ROSC to analyze several associations. In Cohort 1, these parameters included: (i) time to AC-ECoG SD onset during ACA; (ii) time to AC-ECoG RP onset post-ROSC; (iii) area under curve (AUC) of MAP from start of asphyxia to SD onset; (iv) AUC of MAP from ROSC to RP onset.

In Cohort 3, the parameter set was comprised largely of quantities obtained from the optical measurements. In this cohort, SD and RP onset were most reliably measured via spatial propagation of tissue scattering. The parameters extracted from Cohort 3 included: (i) spreading ischemia and spreading edema onsets during ACA; (ii) RP onset via scattering wave post-ROSC; (iii) percentage change in scattering during SD; (iv) AUC of CBF curve from start of asphyxia to SD onset; (v) AUC of CBF curve / AUC of CMRO2 curve from start of asphyxia to SD onset; (vi) CBF/CMRO2 at SD onset; (vii) AUC of MAP, CBF, and CMRO2 from ROSC to RP onset.

#### Neurological Outcome Categorization

The high vs low NDS cutoff for representative rats from Cohort 1 was defined by the median NDS of 49. In Cohort 3, rats with median AC-ECoG IQ > 0.75 from 90-100 mins post-ROSC were defined as high IQ, and median AC-ECoG IQ < 0.75 at 90-100 mins post-ROSC as low IQ.

#### Statistical Models and Tests

Statistical testing was performed with R (v.1.1.456) (RStudio, Boston, MA) and MATLAB. Unsupervised hierarchical clustering of Pearson (Cohort 1) and Spearman (Cohort 3) correlations were performed for correlation matrices with the ggcorrplot package (v0.1.2) of R (Fig. S1). To develop linear regression models for Cohort 1, the leaps package (v3.0), a regression subset selection package, was utilized. For analyses involving Cohort 3, 5- and 7-min ACA rats were combined, as asphyxial duration was not a significant covariate in the analysis. Wilcoxon rank-sum tests were performed for analyses comparing the high and low IQ groups in Cohort 3.

Using linear regression models generated for Cohort 1, receiver operating characteristic (ROC) curves were generated by varying the NDS threshold for good vs poor neurological outcome. Optimal sensitivity and specificity were determined using Youden’s J statistic.

## Results

### Multimodal Detection of Spreading Depolarization (SD) during ACA

In our hypoxic-ischemic model of CA, AC-ECoG silencing, also described as non-spreading depression,^16^ was induced within 1 min of the start of asphyxia (Fig. 2A,C). SD was detected at approximately 1.9-2.5 min. For Cohort 1, an AC-ECoG marker for SD onset was determined manually by the 2^nd^ negative deflection on the 1-Hz LP ECoG during the period of electrocerebral silence (Fig. 2A-B). The 2^nd^ negative deflection was often prominent and thus utilized to mark SD onset, because the initial deflections associated with SD onset were not readily apparent across different cohorts of rats. AC-ECoG determination of SD onset was verified with DC potential recordings in Cohort 2 (Fig. 2A). In Cohort 3, SD was not visible on AC-ECoG, possibly due to electrical noise introduced by optical instruments, along with prolonged surgeries inducing higher stress. Instead, SD onset for Cohort 3 was marked by the onset of the scattering wave (Fig. 2D), indicative of cytotoxic edema. SD also coincided with a steady increase in tissue oxygenation (StO2) (Fig. 2C).

**Figure 2.**
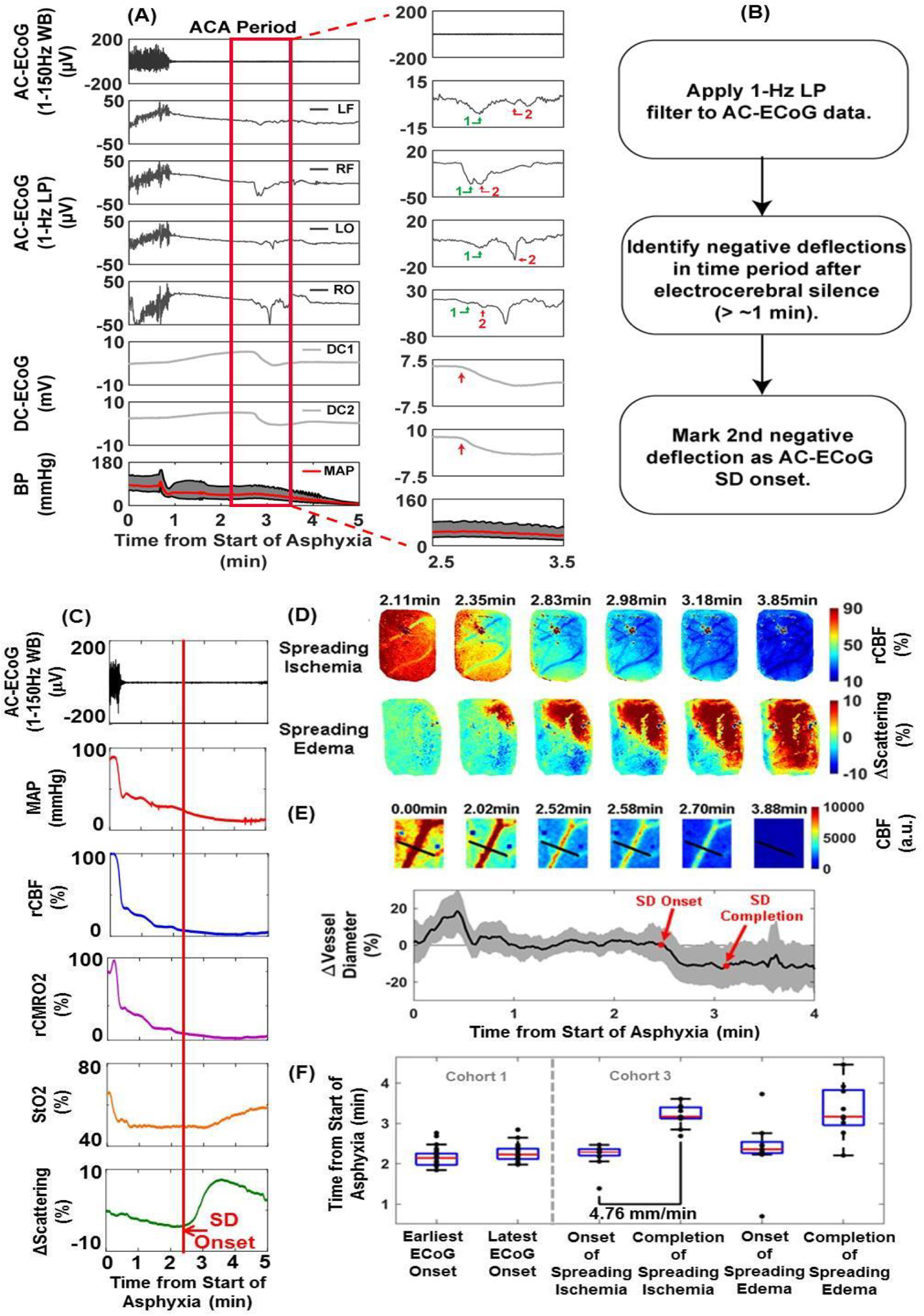
Spatiotemporal features of SD during ACA are detected and characterized with multiple measurement modalities. (A-B) As confirmed by DC-ECoG in Cohort 2, SD from ACA appears as an ultra-slow wave in 1-Hz low-pass (LP) AC-ECoG in Cohorts 1 and 2 for 8-min ACA. This feature is not visible in 1-150 Hz wide-band (WB) AC-ECoG. SD occurs after electrocerebral silence. SD onset for each channel (LF = left frontal; RF = right frontal; LO = left occipital; RO = right occipital) was marked manually on 1-Hz LP AC-ECoG traces in Cohort 1 by the 2^nd^ negative deflection. The arrows and numbers denote the 1^st^ and 2^nd^ negative deflections. (C) A gradual decrease and subsequent rapid increase in tissue scattering, attributed to cytotoxic edema and dendritic beading, is observed during SD. In addition, an upward deflection in tissue oxygenation (StO_2_), attributed to cessation of metabolism, is observed immediately following SD. Onset of SD, as defined by onset of spatial propagation of tissue scattering change in optical imaging window, is denoted by the gray vertical line. rCBF and rCMRO2 denote changes relative to pre-asphyxia baseline in CBF and CMRO2. (D-E) Hallmarks of SD measured with optical imaging include spreading ischemia (spatially-propagating CBF wave) and spreading edema (spatially-propagating tissue scattering wave). SD is confirmed by measuring vasoconstriction during the spreading waves. (F) Consistent with previous reports^60^ of multifocal SD onsets, earliest and latest SD onsets across the 4 AC-ECoG channels in Cohort 1 were separated by 7.2 ± 7 sec (x□ ± σ). The speed of spreading ischemia, 4.76 ± 1.10 mm/min (x□ ± σ), was also consistent with previous reports.^61^

### Spatiotemporal SD-Related Features during ACA

In Cohort 3, spreading ischemia was observed to travel from lateral to medial within the craniectomy (Fig. 2D-F), in agreement with SD directionality from previous publications.^29,62^ The detected vasoconstriction (Fig. 2E) and measured speed of spreading ischemia (∼4.75 mm/min; Fig. 2F) are in agreement with previously-reported SD characteristics.^61^ SD onset did not always occur simultaneously across the cortex in Cohort 1 (Fig. 2F). The largest observed range of onset times across the 4 channels was 28 sec.

### Earlier Depolarization and Smaller Increases in Scattering are Associated with Better Neurological Outcome

Earlier SD onset in Cohort 1 was associated with better neurological outcome (Fig. 3A-B). The earliest onset among the 4 channels provided the strongest correlation with NDS, with mean and latest channel onsets providing progressively weaker correlations (Fig. S2). Earliest SD onset was used for further analysis. In Cohort 3, earlier SD onset trended towards higher AC-ECoG IQ (Fig. S1). Notably, a smaller increase in tissue scattering during SD was associated with higher AC-ECoG IQ post-ROSC (Figs. 3C-E).

**Figure 3.**
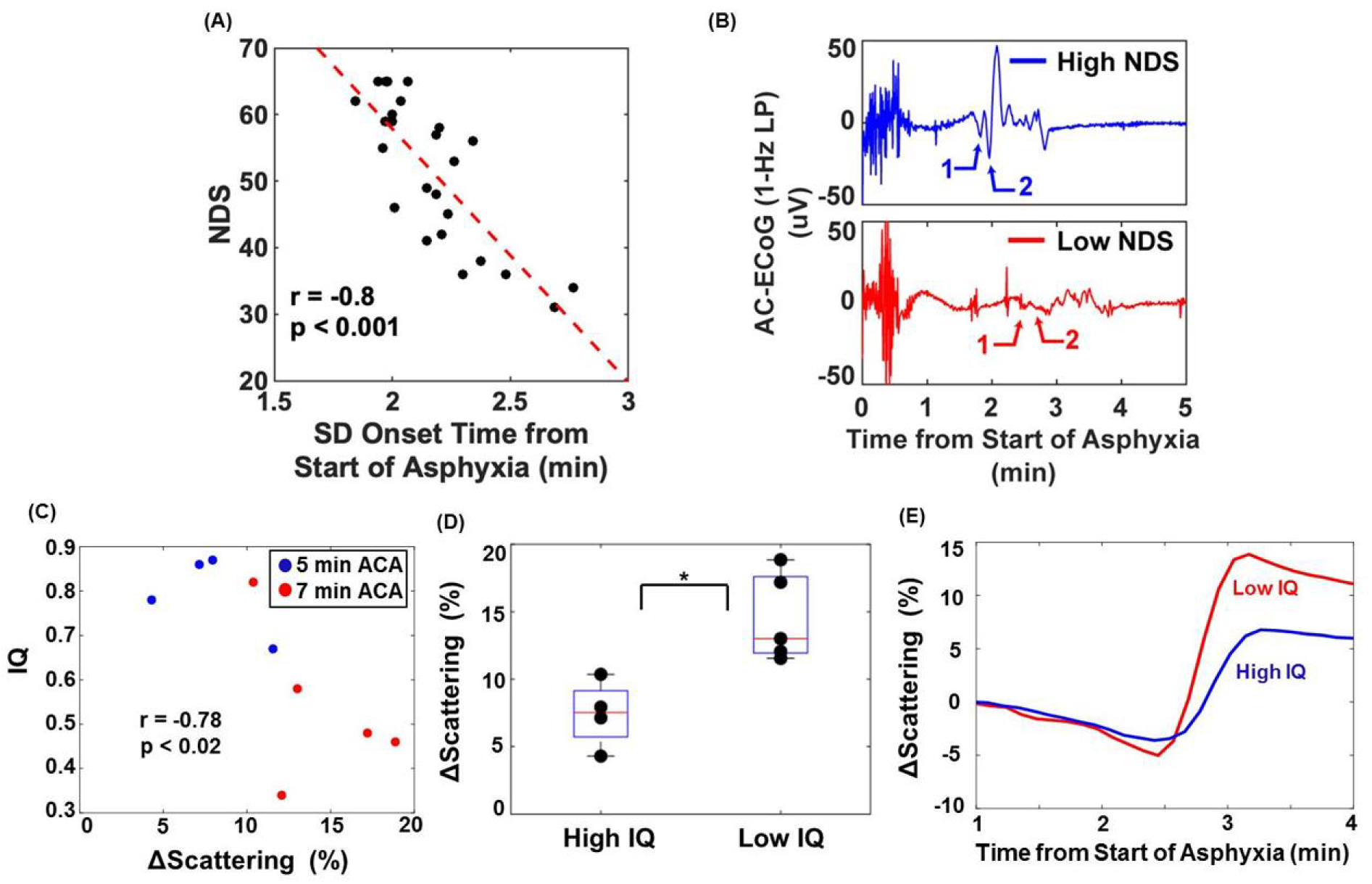
Delayed SD onset and increased cytotoxic edema are associated with poor outcome. (A) Earlier LP AC-ECoG SD onset is associated with better neurological outcome (NDS) in Cohort 1. (B) Representative 1-Hz LP AC-ECoG tracings show earlier SD onset for a rat with good NDS (NDS ≥ 49) and later onset for a rat with poor NDS (NDS < 49). (C) Smaller increases in tissue scattering during SD are associated with higher AC-ECoG IQ post-ROSC. (D) Rats with low IQ exhibit larger increases in tissue scattering. (E) Representative tissue scattering tracings show greater increase in tissue scattering for a rat with low IQ.

### Algorithmic Determination of AC-ECoG SD Onset

To verify manual detection of SD onset for Cohort 1, an automated algorithm was developed to similarly determine the 2nd negative deflection for SD onset (Fig. S3). The automated SD onsets confirmed the manual onsets (r = 0.9) and the association between SD onset and NDS (r = −0.68, p < 0.001).

### Greater Global Perfusion, Cerebral Perfusion, and Higher CBF/Metabolism ratio until SD onset is Associated with Worse Neurological Outcome

Higher total peripheral blood pressure (MAP) and CBF, from start of asphyxia to SD onset, were associated with worse neurological outcome (Fig. 4A, Fig. S1). In Cohort 1, higher average blood flow (i.e., mean MAP) was also associated with lower NDS (Fig. S1A). Additionally, in Cohort 3, low IQ rats had higher CBF (Figs. C, D) and higher flow-metabolism ratio (Figs. 4E-H), suggesting excess cerebral perfusion during ACA.

**Figure 4.**
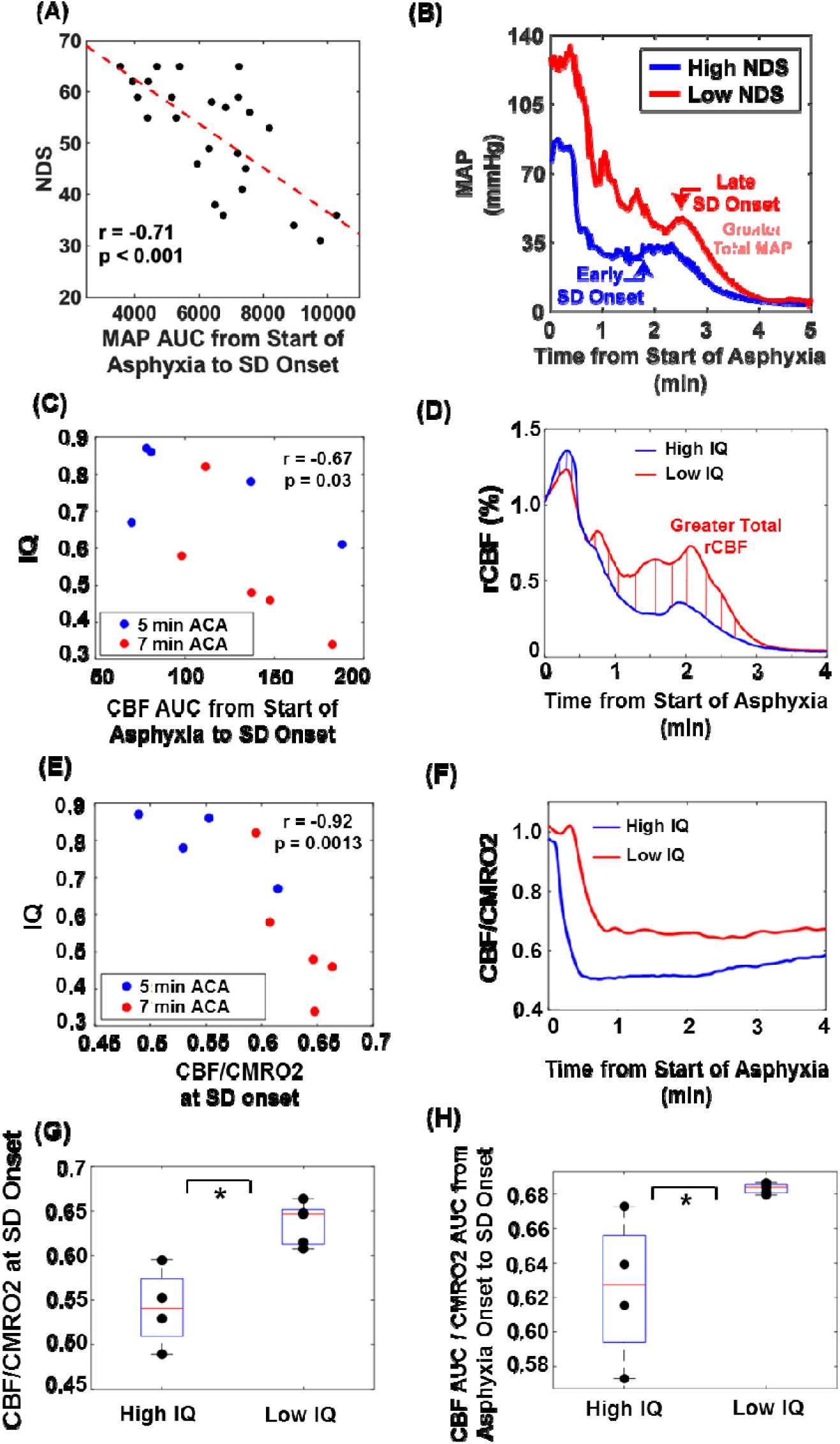
Greater perfusion and higher cerebral perfusion-metabolism ratio up to SD onset correlate with worse outcome. (A-B) Reduced total (AUC; area under the curve) mean arterial pressure (MAP) and (C-D) cerebral blood flow (CBF) during entry into CA (asphyxia up to SD onset) are associated with better neurological recovery (24-hr NDS, (A); ECoG IQ 90 min post-ROSC, (C)). (E-G) Lower cerebral perfusion/metabolism ratio (CBF/CMRO2) at SD onset is associated with higher IQ. (H) Rats with high IQ also have lower ratio of total cerebral perfusion (CBF AUC) to total cerebral metabolism (CMRO2 AUC) over the period from start of asphyxia to SD onset.

### Multimodal Detection of Repolarization Following Resuscitation

RP onset was identified on low-pass AC-ECoG by utilizing MAP (Fig. 5A-B). In Cohort 2, it was observed that RP on the DC channels, marked by a rise in the DC potential, invariably initiated during the downslope of the initial MAP spike following ROSC. RP onset was marked by a positive deflection in the low-pass AC-ECoG during the initial period of post-ROSC MAP decrease. Due to artifact or inability to determine RP onset, 5 (out of 27) rats of Cohort 1 were excluded from further RP onset analysis. AC-ECoG determination of RP onset was more difficult to determine than SD onset (Fig. S4). RP onset for Cohort 3 was determined by visual inspection of the tissue scattering maps, which showed complex spatiotemporal dynamics during this time period (Fig. 5D).

**Figure 5.**
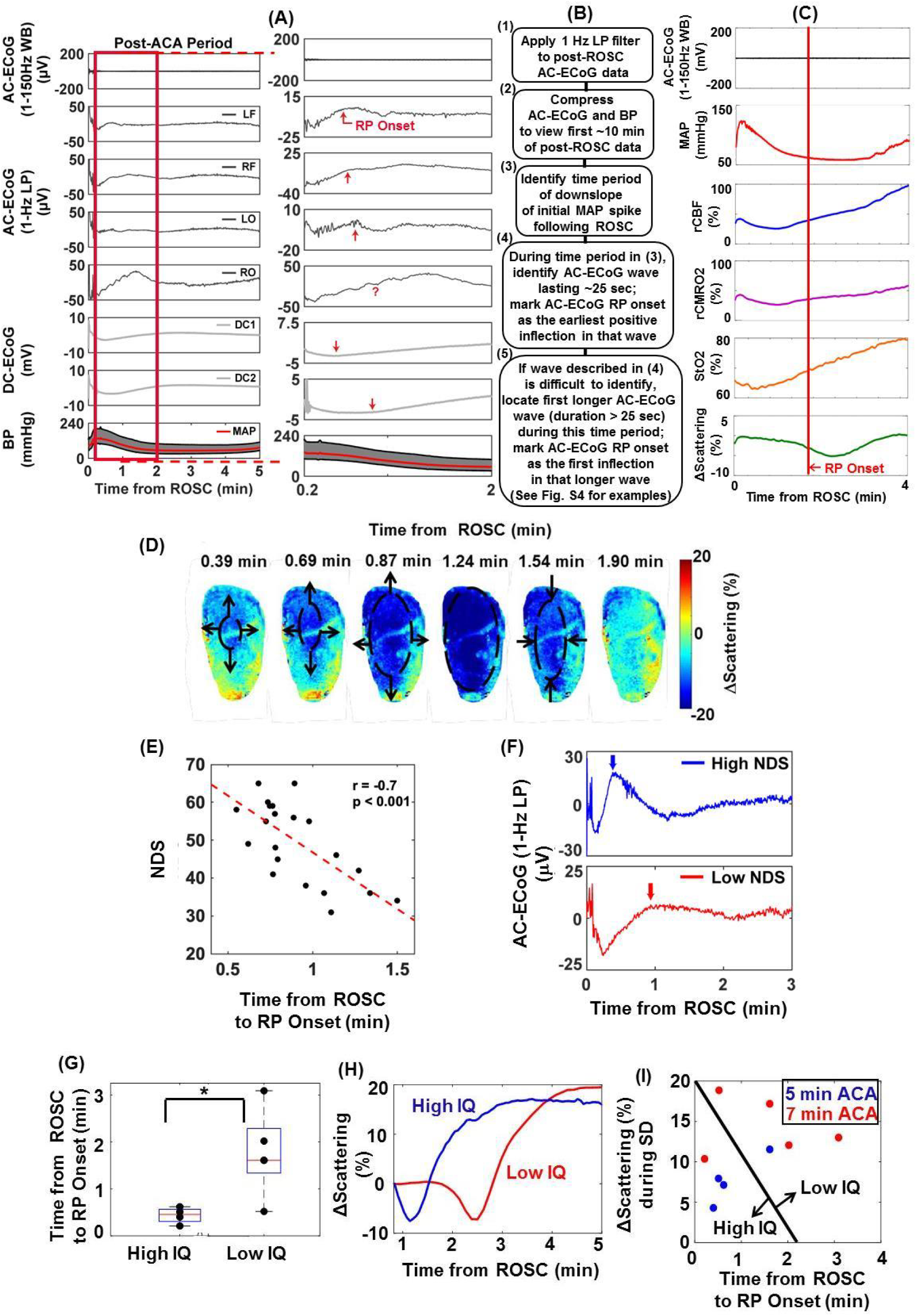
Repolarization (RP) following ROSC is detected with multiple measurement modalities, and earlier RP onset is associated with better outcome. (A-B) Post-ROSC RP is not visible in 1-150 Hz WB AC-ECoG, but large waves are visible in 1-Hz LP AC-ECoG. RP onset was discovered to occur during the downslope of the spike in post-ROSC BP. RP onset is marked by a large, slow positive deflection on LP AC-ECoG during the BP downslope, approximately corresponding to an inflection point on the DC potential. (C) An inflection point in tissue scattering (attributed to neuronal RP) is observed during the same time period as the wave in (A), even though no such feature is observed in cerebral blood flow (rCBF), cerebral metabolic rate of oxygen (rCMRO2), or tissue oxygenation (StO2). Onset of RP, as defined by onset of spatial propagation of tissue scattering change in optical imaging window, is denoted by the red vertical line. (D) Spatiotemporal changes in rat brain scattering coefficient within the first 2 min post-ROSC provide a potential optical biomarker of spreading repolarization. (E-F) Earlier RP onset in Cohort 1 is associated with better neurological recovery (NDS). Arrows indicate RP onset. (G-H) For Cohort 3, RP onset was determined by start of the scattering wave, and high IQ rats repolarized significantly earlier than low IQ rats. (I) When RP onset was plotted against the percentage change in scattering during SD, high and low IQ rats formed separate clusters, and rats with smaller scattering increases during SD tended to repolarize earlier.

### Earlier Repolarization is Associated with Better Neurological Outcome

Earlier ECoG RP onset in Cohort 1 was associated with better neurological recovery (Fig. 5E-F). Similar to SD onset, the earliest RP onset among the 4 channels exhibited the strongest correlation with NDS (Fig. S4), so earliest onset was again utilized for further analysis.

High NDS rats had significantly earlier RP onset than low NDS rats (Fig. 5E-F). In Cohort 3, high IQ rats had significantly earlier RP onset times than low IQ rats, as measured with tissue scattering (Fig. 5G-H). Interestingly, when RP onset was plotted against change in scattering during SD, high and low IQ rats separated into distinct clusters, with high IQ rats tending to have earlier RP onset and smaller increases in scattering during SD (Fig. 5I).

### Total Depolarization Duration from SD Onset to RP Onset Is Not Associated with Neurological Outcome

Since the depolarization period is commonly believed to be injurious,^63,64^ we examined the total combined duration of depolarization pre- and post-CPR. There was no association between combined depolarization duration and NDS, likely due to earlier depolarizing rats also repolarizing earlier (Fig. S6).

### Higher Global Perfusion, Cerebral Perfusion, and Cerebral Metabolism from ROSC to RP Onset Are Associated with Worse Neurological Outcome

More total peripheral (global) blood flow, CBF, and cerebral metabolism from ROSC to RP onset trended toward worse neurological outcome (Fig. 6, Fig. S1). Rats in the low IQ group had significantly greater peripheral and cerebral perfusion and metabolism during this period than rats in the high IQ group (Fig. 6B-F). Average MAP from ROSC to RP onset was not correlated with NDS (Fig. S1A), which may indicate that the aforementioned RP relationships are primarily dictated by duration of time between ROSC and RP onset (Fig. 5E-G).

**Figure 6.**
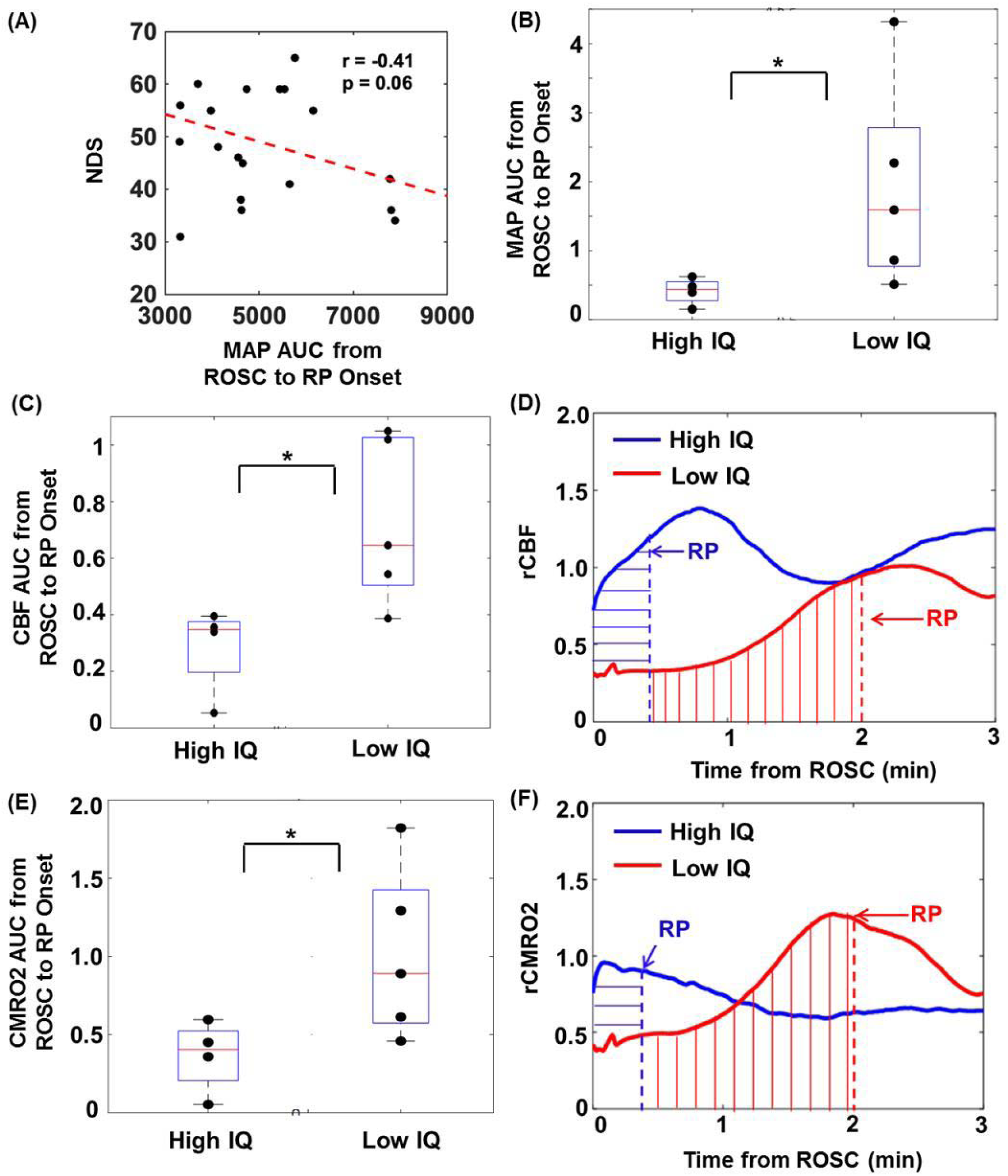
Greater reperfusion and cerebral metabolism from ROSC to RP onset are associated with worse outcome. (A) In Cohort 1, more total MAP, from ROSC until RP onset, trends toward worse neurological outcome (NDS). (B) In Cohort 3, high IQ rats have significantly less total MAP from ROSC until RP. (C-F) High IQ rats also have significantly less cerebral reperfusion and metabolism from ROSC until RP.

### SD and RP Onsets Predict Neurological Outcome

13 parameters were tested for inclusion in a linear regression model to predict NDS in Cohort 1 (Fig. S1A). SD onset (Fig. 7(A)), RP onset (Fig. S7(A, C)), and baseline glucose (Fig. S7(A), Fig. S8(C)) had the strongest correlations with NDS. SD onset prediction of NDS was significantly improved by addition of RP onset (p=0.018; Fig. 7). RP onset prediction of NDS was significantly improved by addition of baseline glucose (p=0.037; Fig. S7). Using both SD onset and RP onset in the model provided the most accurate predictions of NDS (Fig. 7).

**Figure 7.**
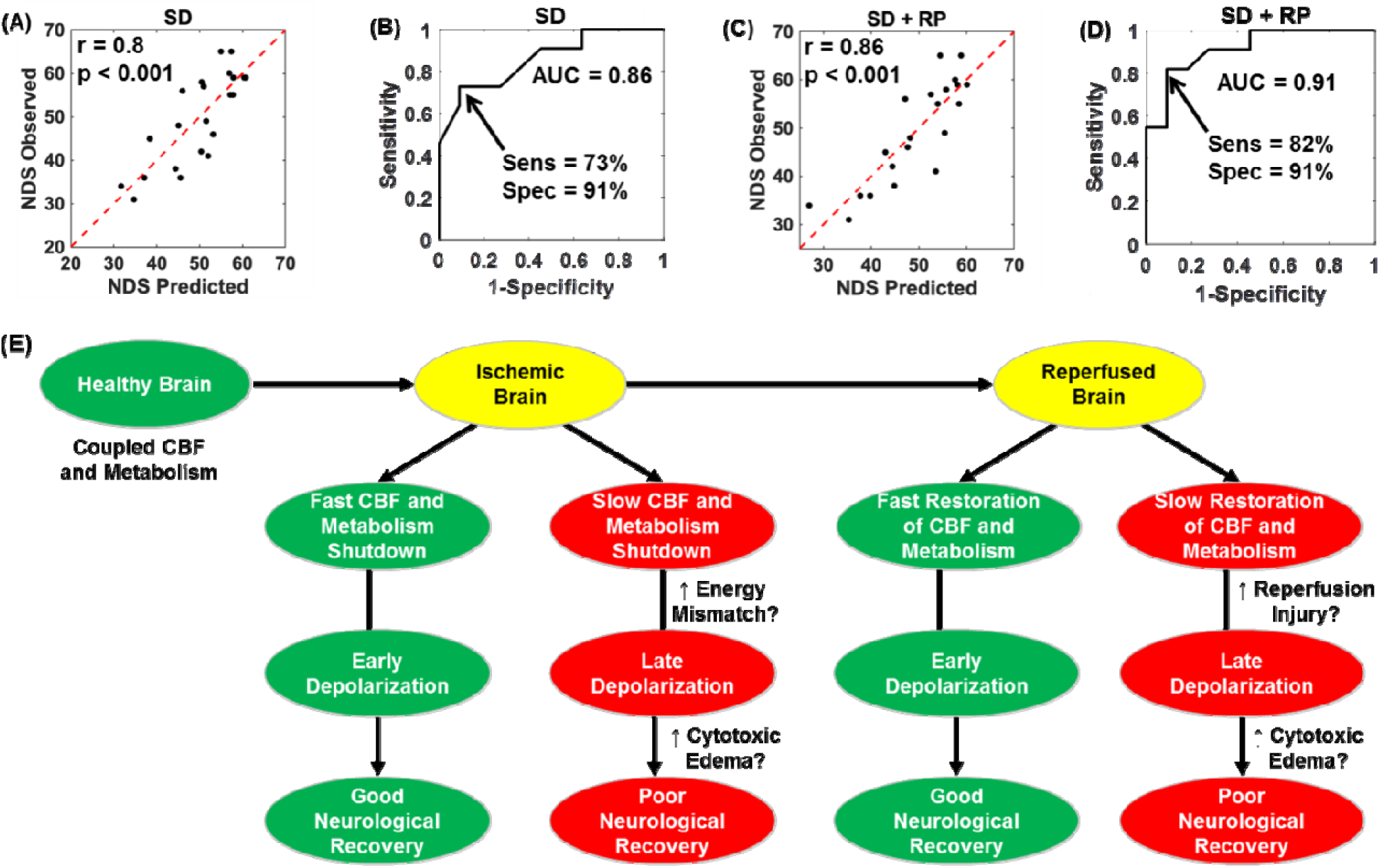
Neurological outcome prediction model with SD and RP onsets. Out of 13 variables for Cohort 1, SD onset (A-B) predicted NDS with greatest accuracy in models using 1 predictor variable (r = 0.8; ROC AUC = 0.86). Optimal sensitivity and specificity, as determined by Youden’s J statistic, were 73% and 91%, respectively. (C-D) Adding RP onset as a second predictor variable improved the accuracy of the model (r = 0.86; ROC AUC = 0.91). Optimal sensitivity and specificity, as determined by Youden’s J statistic, were 82% and 91%, respectively. (E) Flow chart outlining associations between SD, RP, and neurological outcome. Early SD and RP are associated with good neurological outcome, while delayed SD and RP are associated with poor neurological outcome.

The 4 models were further utilized to predict high vs low NDS. SD onset and RP onset alone correctly predicted 18/22 rat outcomes (81.82% accuracy), while SD onset plus RP onset and RP onset plus baseline glucose correctly predicted 20/22 outcomes (Fig 7; 90.91% accuracy).

## Discussion

In this report, we investigate the relationship between the timing of SD dynamics during CA and neurological outcome 90 min and 24 hrs after CA+CPR. To our knowledge, we provide the first demonstration that earlier SD during CA is prognostic of better neurological outcome after resuscitation (r = −0.80, p < 0.001). We verify that this phenomenon has the characteristics of SD by measuring a DC potential shift concomitant with the observed AC-ECoG ultra-slow wave. The SD onsets detected with our manual algorithm were confirmed by an automated algorithm (r = 0.9), which supported the association between SD onset and NDS (r = −0.68, p < 0.001). Additional verification is provided by optically-measuring spreading ischemia (via CBF) and edema (via tissue scattering) with similar onset time as the ultra-slow ECoG wave. Tissue scattering changes are indicative of cytotoxic edema and neuronal injury,^1,24,40,65^ and the magnitude of the scattering increase during SD correlated negatively (r = −0.78, p < 0.02) with short-term neurological recovery (ECoG IQ 90 min post-ROSC). This finding supports the hypothesis that the scattering increase during SD may lead to poor short-term outcome measures and may be attributed largely to dendritic beading, an indicator of the amount of neuronal damage.^40,66^

Upon resuscitation, we detect a corresponding RP with AC-ECoG, DC potential, and tissue scattering. Earlier onset time of this RP also correlated with better neurological recovery (r = −0.70, p < 0.001). To our knowledge, this is the first study to characterize post-CA RP using AC-ECoG and optical imaging and correlate RP-related parameters with neurological outcome. Combining SD and RP onset times in a multiple linear regression model enabled prediction of 24-hr NDS with a sensitivity, specificity, and ROC AUC of 82%, 91%, and 0.91, respectively. In addition, the combination of RP onset and change in tissue scattering during SD separated low ECoG IQ rats and high ECoG IQ rats into separate clusters. Thus, less tissue scattering during CA-induced SD was linked with faster RP post-ROSC, which led to higher ECoG IQ 90 minutes post-ROSC. In short, the various findings related to SD and RP features reported in this study may provide ultra-early prognostic markers of neurological outcome for CA patients.

### Earlier SD during CA may be neuroprotective

It has been posited in multiple recent studies that SD is harmful to the brain,^12,24,67,68^ although this hypothesis is often focused on prolonged SD or repetitive SD events.^68,69^ However, several studies have suggested that mitochondrial processes that are important in SD can signal pathways to protect neurons and endothelium.^70,71^ Indeed, multiple studies have shown beneficial properties of SD, including preconditioning, upregulation of growth factors, and neurogenesisis.^72–77^ In addition, previous studies in insects^78–81^ and rodents^72,73,82,83^ suggest that SD may play a neuroprotective role during ischemia.^1,84^ It is possible that this protection is enabled by a “neuronal silencing” mechanism associated with SD^85^ wherein metabolism is quickly shut down, possibly to conserve residual energy and minimize production of reactive oxygen species, potentially leading to an improved survival and outcome in reversible processes.^81,86^ In this report, we demonstrate that earlier SD may be neuroprotective in a model of transient global cerebral ischemia and reperfusion. Furthermore, we observe correlations between baseline glucose and SD and RP onsets (Fig. S8). Rats with higher levels of baseline glucose had later onsets of SD and RP, as well as lower 24-hr NDS. This result suggests that depletion of energy stores in the brain trigger SD, and that if this process happens earlier during entry into CA, it may help lead to better neurological recovery post-CPR.

### Early SD reduces periods of mismatch between CBF and brain metabolism during CA

In Fig. 7(E), we propose a mechanism by which early SD and RP may be neuroprotective in CA. In the presence of anoxia leading to global cerebral ischemia (e.g. CA), reduced oxygen metabolism and gas exchange in the brain during periods of low flow CBF may be related to the formation of harmful reactive oxygen species.^87–89^ Evidence for this hypothesis can be found in several different measurement parameters from our data. First, metrics related to total perfusion from start of ischemia/anoxia to onset of SD (AUC of MAP and CBF) are lower in animals that recovered better post-CPR. Second, the ratio of CBF to CMRO2 (an indicator of the cerebral perfusion-metabolism mismatch described previously^38^) was lower at SD in animals that recovered better after CPR. This may suggest that continued perfusion during hypo-metabolism and/or anoxia in the brain is detrimental. Recently, it has been shown that formation of these reactive oxygen species during cerebral ischemia may be mitigated through manipulation of glucose metabolism.^87^ Future experiments utilizing our multimodal platform to test these concepts and maneuvers on cerebral perfusion and metabolism throughout the dynamic periods of ischemia and reperfusion can help to uncover the underlying mechanisms of how flow-metabolism dynamics during CA-induced SD affect outcome.

### Early RP may mitigate reperfusion injury following resuscitation

Following ROSC, we hypothesize that a similar phenomenon occurs. During the transient period from ROSC to RP, reperfusion occurs without sufficient corresponding recovery of cerebral metabolic activity. Therefore, the brain may be unable to consume enough of the newly-available oxygen. Specifically, animals with less total CBF prior to RP have better outcome. As during entry into CA, a flow-metabolism mismatch may lead to reperfusion injury, potentially by formation of reactive oxygen species,^90^ worsening neurological recovery. This theory is supported by our data that rats with worse neurological recovery had greater total perfusion (MAP AUC, CBF AUC) from ROSC to RP.

### Limitations

It is important to acknowledge that this report does not include measurements of cerebral changes and neurological recovery on the molecular, cellular, and histological level. Work in this area is essential to confirm the hypothesis suggested in Fig. 7E. Second, the animals were split into multiple cohorts with slightly different surgical preparation and ECoG implantation schemes to accommodate the multiple data collection modalities. One consequence of this is that NDS was unavailable for animals in the optical imaging cohort, as they were sacrificed after 2 hrs of monitoring following ROSC due to the craniectomy. Therefore, it was necessary to use ECoG IQ 90 min post-ROSC as a surrogate outcome metric for this cohort. This limitation will be addressed in future studies by using a chronic imaging window^91,92^ to enable survival studies out to 1 month post-ROSC. Additionally, the craniectomy was performed over only a 6 mm × 4 mm area of the skull, precluding the entire surface of the brain to be exposed for imaging. As a result, our measurements of SD onset in Cohort 3 was based on where and when the CBF and scattering changes first appeared within the craniectomy region, not the brain as a whole. Furthermore, it is known that DC potential readings can be distorted by changes in CBF and pH,^93^ so it is unclear how much of the DC shift measured in Cohort 2 during SD and RP was caused by electrophysiological changes in the brain, as opposed to being an artifact of a concomitant change in CBF and pH. However, we believe that we were able to address this issue by employing tissue scattering as an independent measure of SD and RP in Cohort 3. Finally, the method by which we identified SD onset and RP onset on the AC-ECoG signal was novel and subjective and would benefit from further validation in other paradigms. However, we also developed an automated algorithm to detect SD onset on the AC-ECoG signal that is highly correlated with the operator-identified method that we utilized.

## Conclusion

In this report, we use a preclinical model mimicking an intensive care unit along with a multimodal monitoring setup including optical imaging and electrophysiology to characterize SD in the brain during CA, RP in the brain following resuscitation, and the relationship between SD/RP parameters and neurological recovery. We demonstrate, for the first time, that earlier SD onset correlates with, and is predictive of, improved neurological outcome (24-hr NDS). Including RP onset into this predictive model provides an additional improvement over using SD alone. We also show that lower values of optically-measured parameters related to CBF, metabolism, flow/metabolism coupling, and cytotoxic edema during SD and RP are correlated with improved neurological recovery. Characterizing SD and RP in this manner demonstrates strong potential to provide (1) ultra-early prognostication of neurological recovery from global ischemia and (2) elucidation of potentially modifiable mechanisms of cerebral response to global ischemia and reperfusion. These metrics should be tested in future studies to inform clinically translatable interventions for CA patients during CA itself or immediately post-resuscitation, enabling patient-specific treatment at early time points that may be critical for optimizing neurological recovery. These findings may also have implications for other acute brain injury conditions, such as focal stroke, subarachnoid hemorrhage, and traumatic brain injury, where SD can occur.

## Acknowledgements

This work was supported by the Arnold and Mabel Beckman Foundation, the Roneet Carmell Memorial Endowment Fund to Y.A., the United States National Institutes of Health (P41EB015890), the National Science Foundation Graduate Research Fellowship Program (DGE-1321846, to C.C.), the National Center for Research Resources and National Center for Advancing Translational Sciences, National Institutes of Health, through the following grants: TL1 TR001415-01 to R. H.W., R21 EB024793 to Y. A., KL2 TR001416 to Y.A. via UL1 TR001414, and a CTSA pilot grant to Y.A. via UL1 TR001414. The content is solely the responsibility of the authors and does not necessarily represent the official views of the NIH.

## Supplementary Materials

**Table S1.** Arterial Blood Gas (ABG) parameters from Cohort 1. Rats (n=27) were divided into good and poor NDS by NDS=49 cutoff. Poor NDS rats had significantly higher blood glucose. Student’s t-test and false discovery rate (FDR) for multiple comparison correction were performed. * denotes *p* < 0.05, ** denotes *p* < 0.01, *** denotes *p* < 0.001.

| Time Period | Parameter | NDS ≥ 49 | NDS < 49 | p-value | FDR |
| --- | --- | --- | --- | --- | --- |
| Pre-ACA | pH | 7.416 ± 0.06 | 7.416 ± 0.59 | 0.9987 | 0.614 |
|  | pCO<sub>2</sub> (mmHg) | 38.93 ± 8.69 | 41.12 ± 8.08 | 0.5224 | 0.482 |
|  | pO<sub>2</sub> (mmHg) | 178.2 ± 32.5 | 163.3 ± 32.4 | 0.2604 | 0.320 |
|  | HCO<sub>3</sub> (mmol/L) | 24.69 ± 2.46 | 26.15 ± 2.74 | 0.1651 | 0.261 |
|  | Na (mmol/L) | 140.1 ± 1.90 | 138.6 ± 1.80 | 0.0534 | 0.197 |
|  | K (mmol/L) | 3.830 ± 0.53 | 3.920 ± 0.43 | 0.6518 | 0.555 |
|  | Ca (mmol/L) | 1.360 ± 0.06 | 1.391 ± 0.45 | 0.1914 | 0.265 |
|  | Glu (mg/dL) | 136.1 ± 18.5 | 171.0 ± 26.4 | **0.0005\*\*\*** | **0.005\*\*** |
|  | Hb (g/dL) | 11.70 ± 0.59 | 12.29 ± 1.41 | 0.1389 | 0.256 |
| Post-ACA | pH | 7.369 ± 0.15 | 7.394 ± 0.13 | 0.6554 | 0.518 |
|  | pCO<sub>2</sub> (mmHg) | 40.16 ± 13.0 | 38.1 ± 120.7 | 0.6979 | 0.515 |
|  | pO<sub>2</sub> (mmHg) | 281.8 ± 183 | 274.5 ± 120.7 | 0.9023 | 0.624 |
|  | HCO<sub>3</sub> (mmol/L) | 22.38 ± 2.94 | 22.29 ± 2.68 | 0.9385 | 0.611 |
|  | Na (mmol/L) | 145.2 ± 1.50 | 143.6 ± 2.30 | 0.0757 | 0.209 |
|  | K (mmol/L) | 3.200 ± 0.40 | 3.000 ± 0.27 | 0.1057 | 0.234 |
|  | Ca (mmol/L) | 1.146 ± 0.07 | 1.175 ± 0.06 | 0.2708 | 0.300 |
|  | Glu (mg/dL) | 119.1 ± 32.7 | 185.2 ± 69.4 | **0.0159\*** | 0.088 |
|  | Hb (g/dL) | 13.52 ± 0.82 | 13.89 ± 1.23 | 0.4146 | 0.417 |

**Figure S1.**
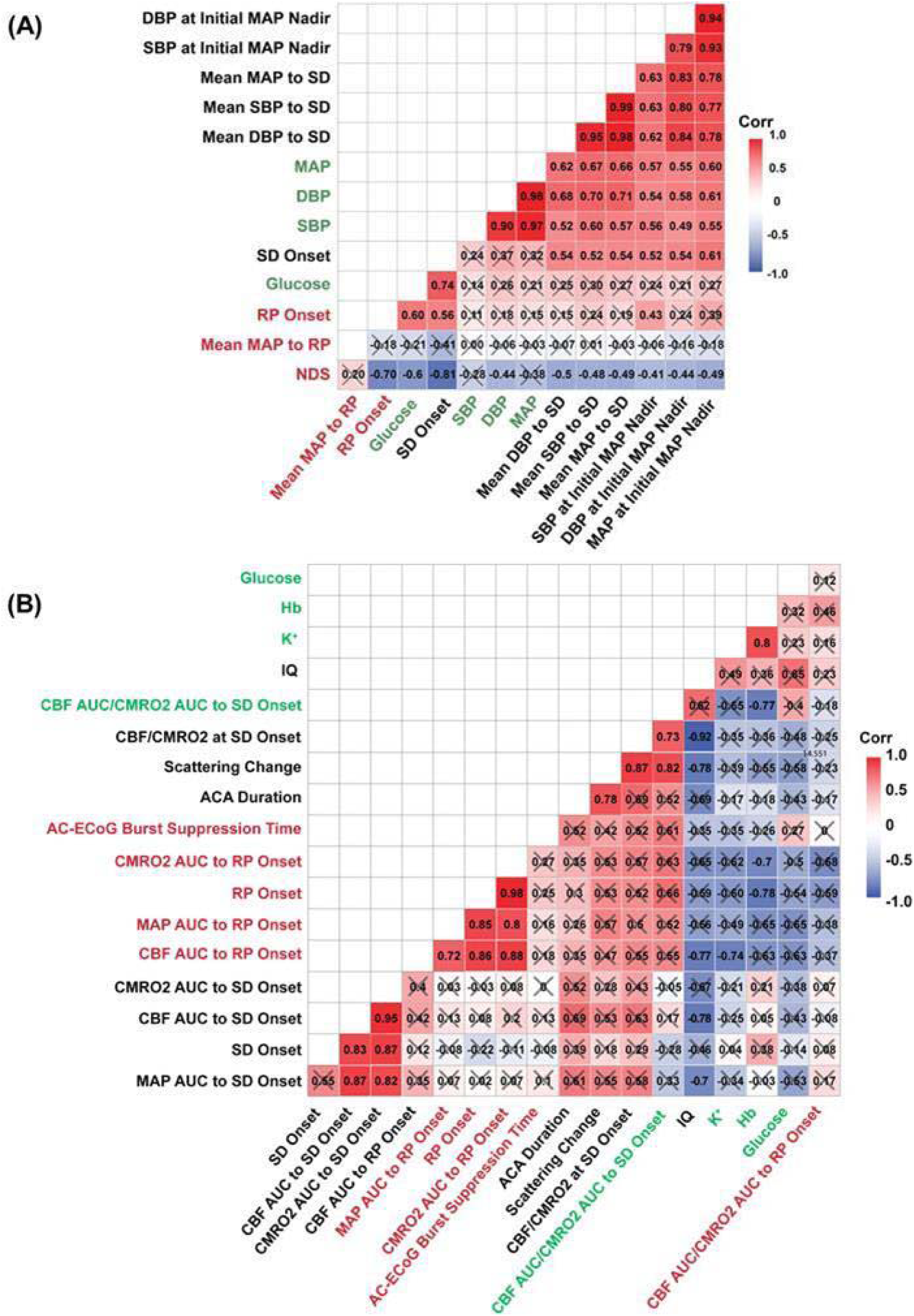
Correlation matrices for Cohort 1 and 3. (A) For Cohort 1, Pearson correlation matrix involving primarily BP, SD, RP, and NDS-related variables. (B) For Cohort 3, Spearman correlation matrix involving ABG, BP, CBF, CMRO2, scattering, and ECoG IQ-related variables. Green indicates pre-asphyxia variables. Red indicates post-asphyxia variables.

**Figure S2.**
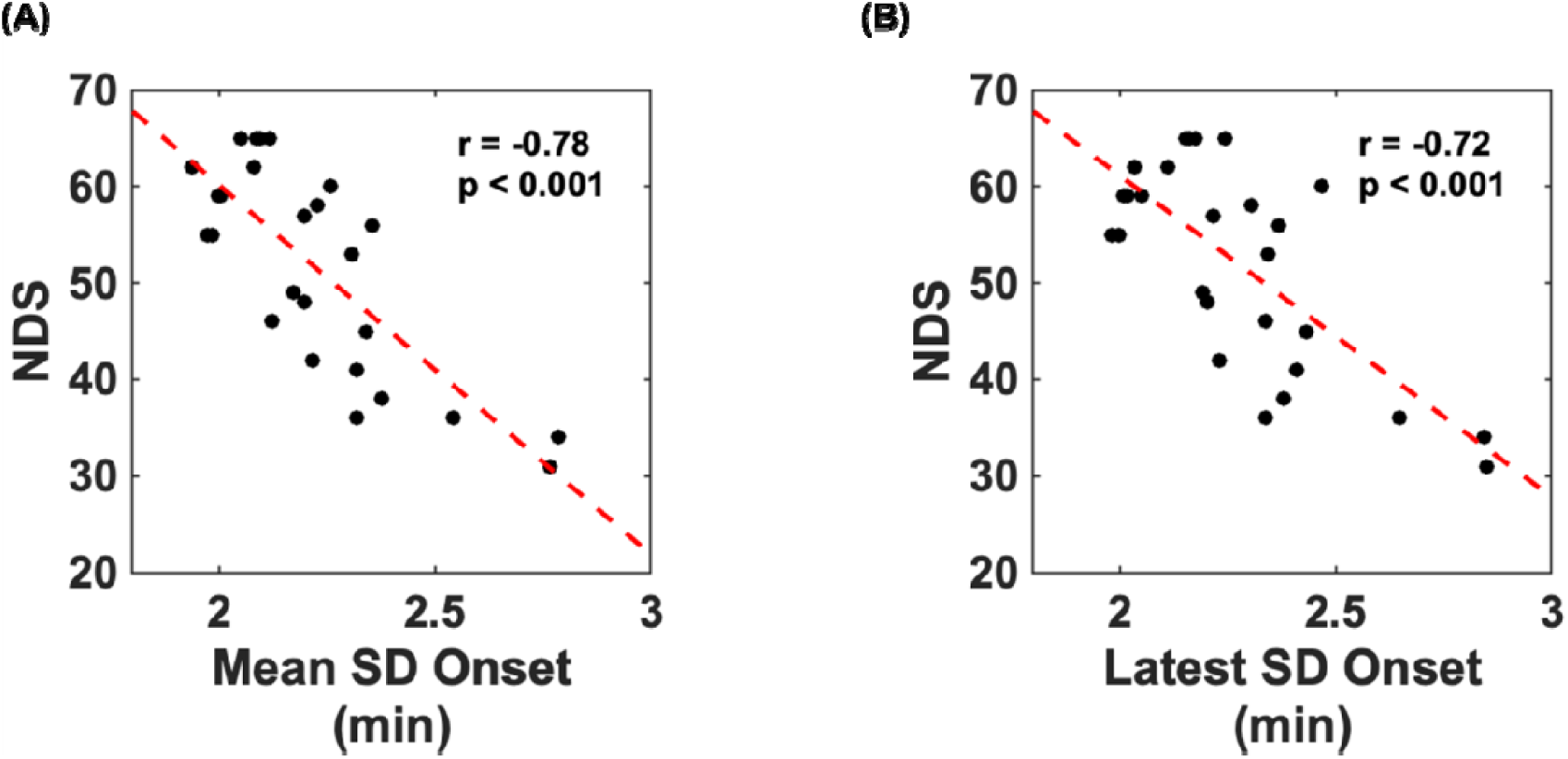
Mean and latest AC-ECoG channel SD onset. In Cohort 1, Mean SD onset of the 4 AC-ECoG channels correlates with NDS (r = −0.78) weaker than earliest channel SD onset (r = −0.8) (Figure 3A). Latest channel SD onset correlates even weaker (r = −0.72).

**Figure S3.**
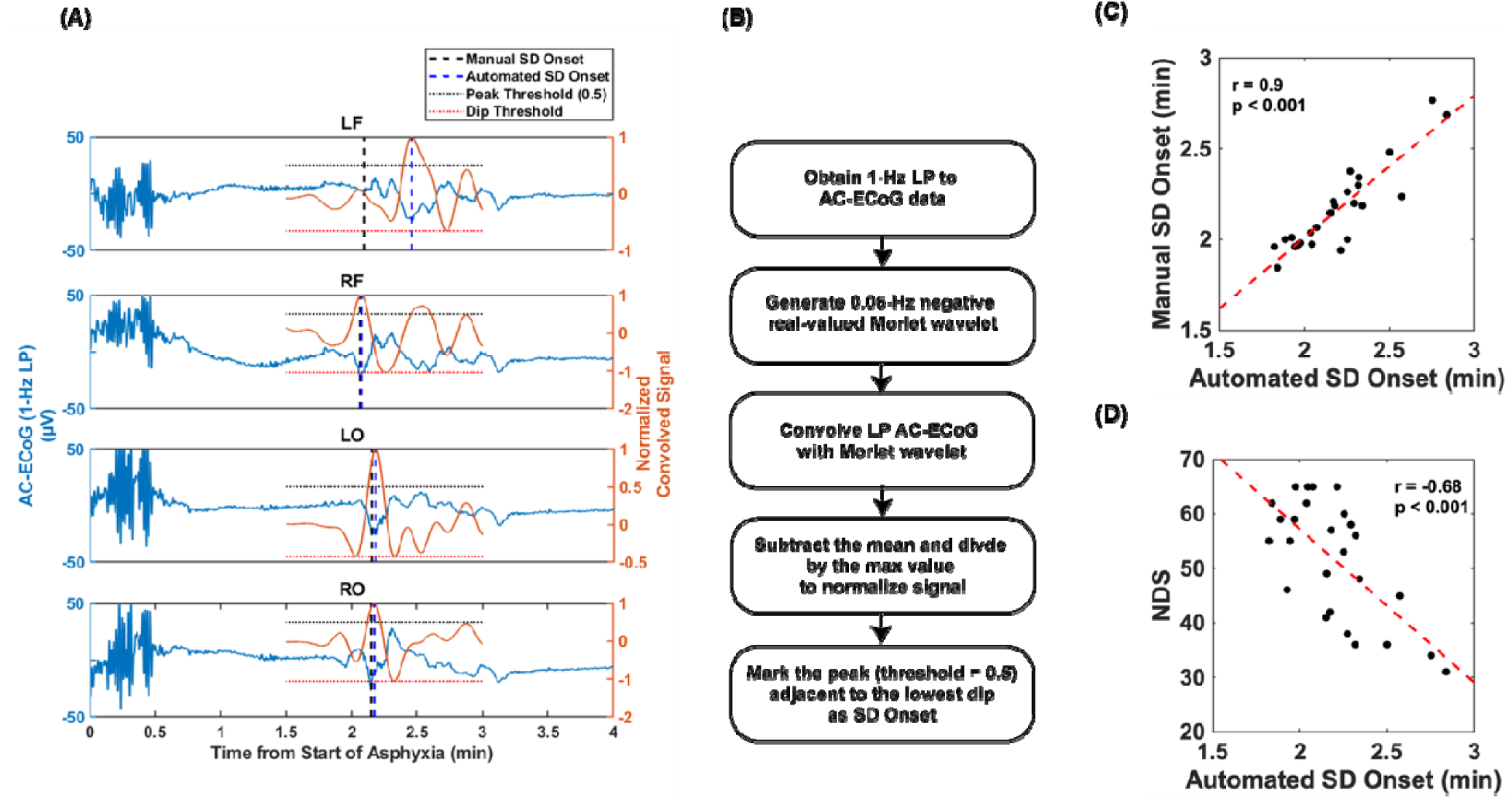
Automated algorithm detection of AC-ECoG SD onset. (A) Representative figure of automated algorithm. (B) Algorithm methodology involving convolution and thresholding for 2^nd^ negative deflection detection. (C) Earliest manual and automated SD onsets correlate strongly (r = 0.9). (D) Automated SD onset correlates with NDS with r = −0.68, while manual SD onset correlates with NDS with r = −0.8 (Figure 3A).

**Figure S4.**
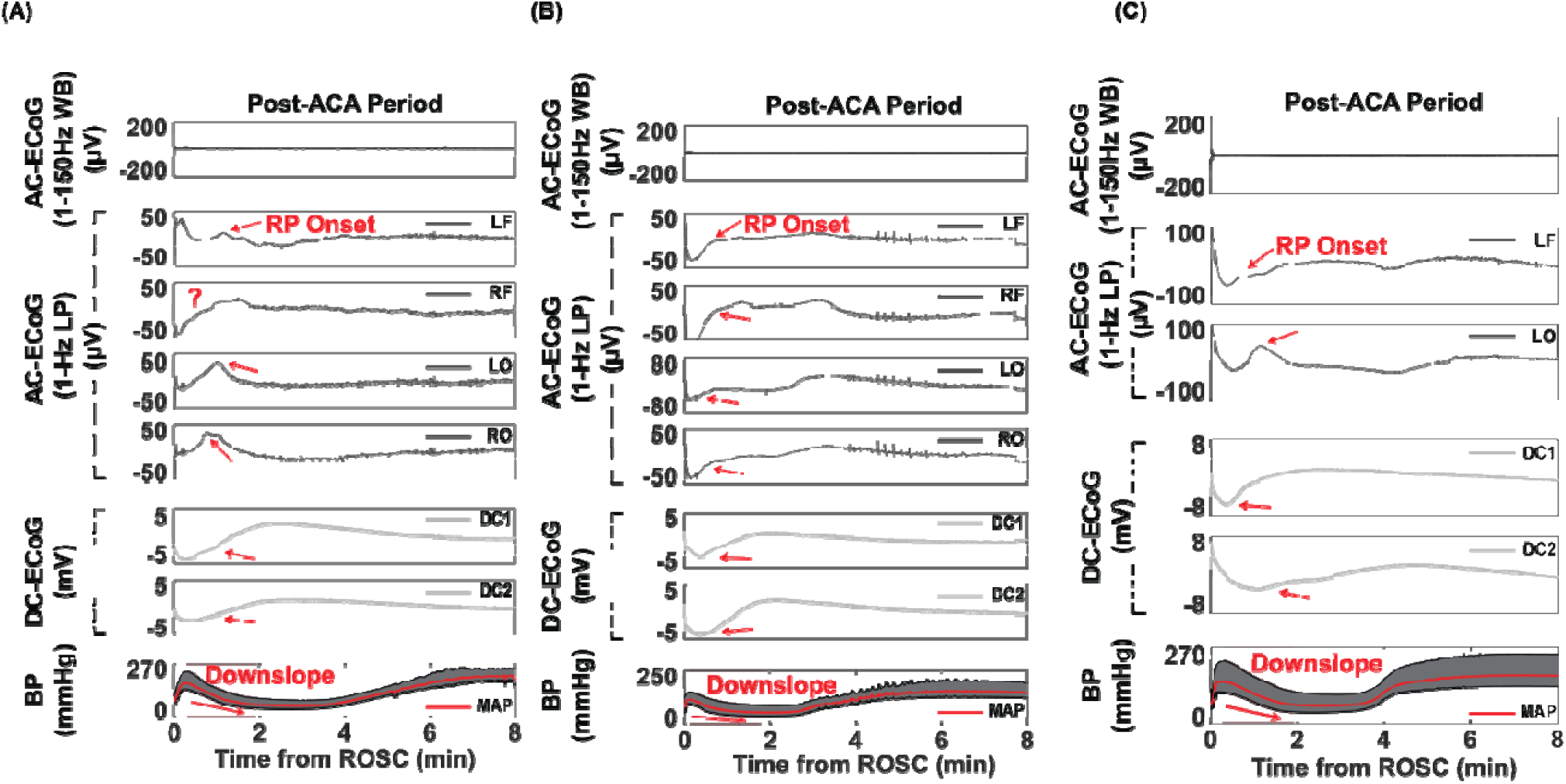
Representative post-ROSC figures from Cohort 2.

**Figure S5.**
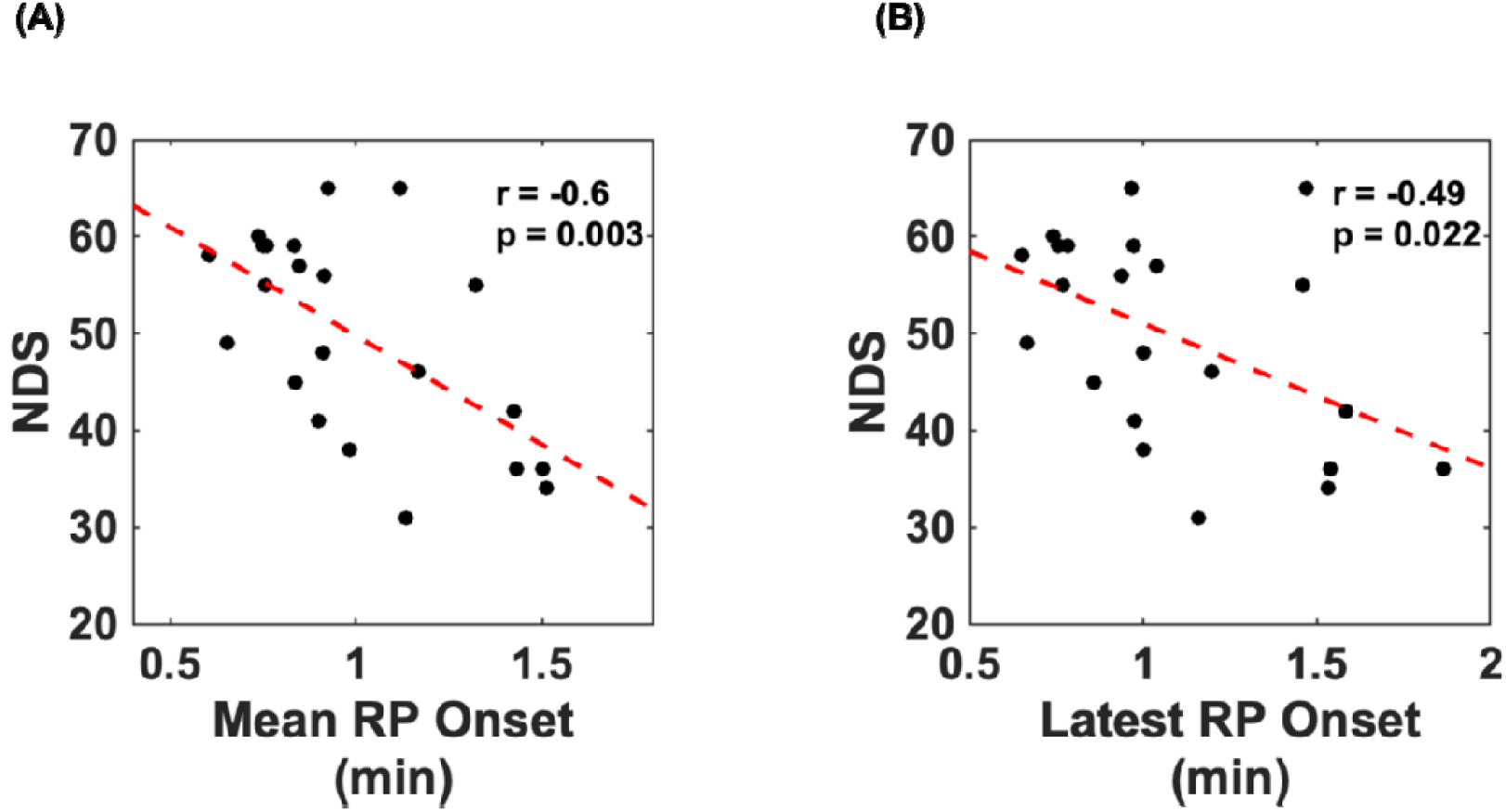
Mean and latest AC-ECoG channel RP onset. In Cohort 1, mean RP onset of the 4 AC-ECoG channels correlates with NDS (r = −0.6) weaker than earliest channel RP onset (r = −0.7) (Figure 5(E)). Latest channel RP onset correlates even weaker (r = −0.49).

**Figure S6.**
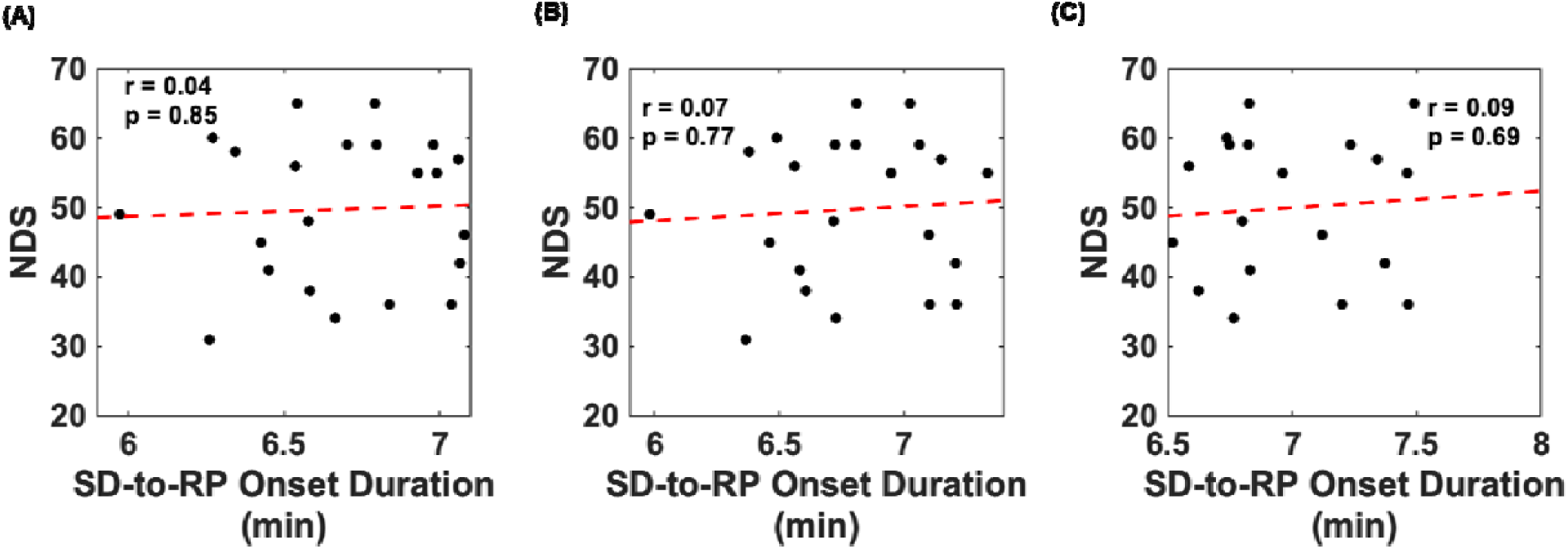
Combined SD duration does not correlate with neurological outcome. In Cohort 1, the time durations from SD onset until start of CPR and from ROSC until RP onset were added to formulate a combined SD duration for each AC-ECoG channel. Although earlier SD onset and earlier RP onset correlated with neurological outcome (Figures 3(A), 5(E)), combined SD duration did not.

**Figure S7.**
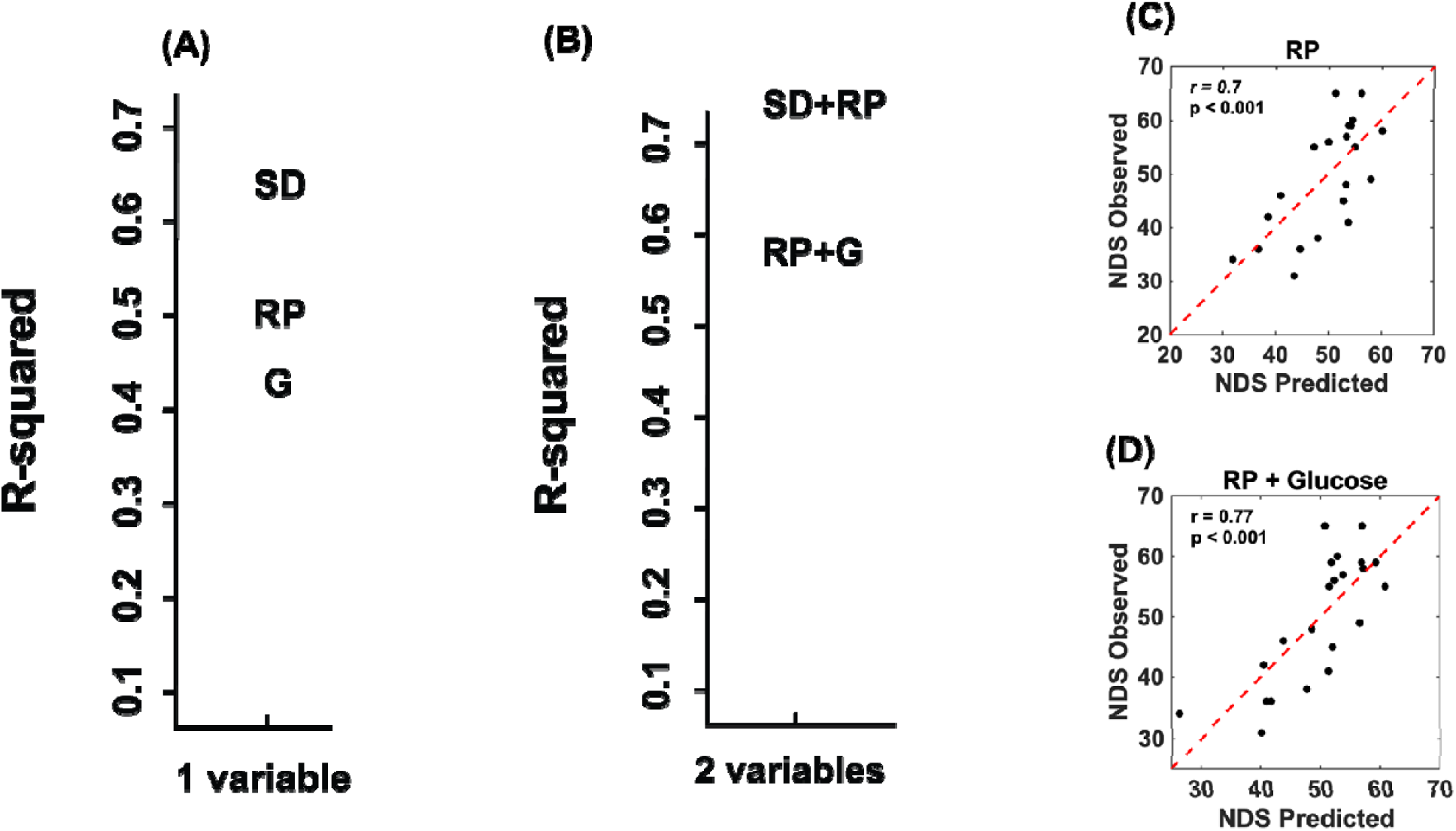
Correlations between linear regression model and NDS. (A) In Cohort 1, SD onset, RP onset, and baseline glucose provided the strongest correlations for 1 predictor models. (B) The RP onset predictor model was significantly improved with addition of baseline glucose. 5 rats were removed from prediction model formulation to match number of RP onset values (n=22). G = baseline glucose. (C) Predicted NDS using RP onset alone correlated with true NDS (r = 0.7). (D) Predicted NDS using RP onset + glucose provided an improved correlation with true NDS (r = 0.77), relative to RP onset alone.

**Figure S8.**
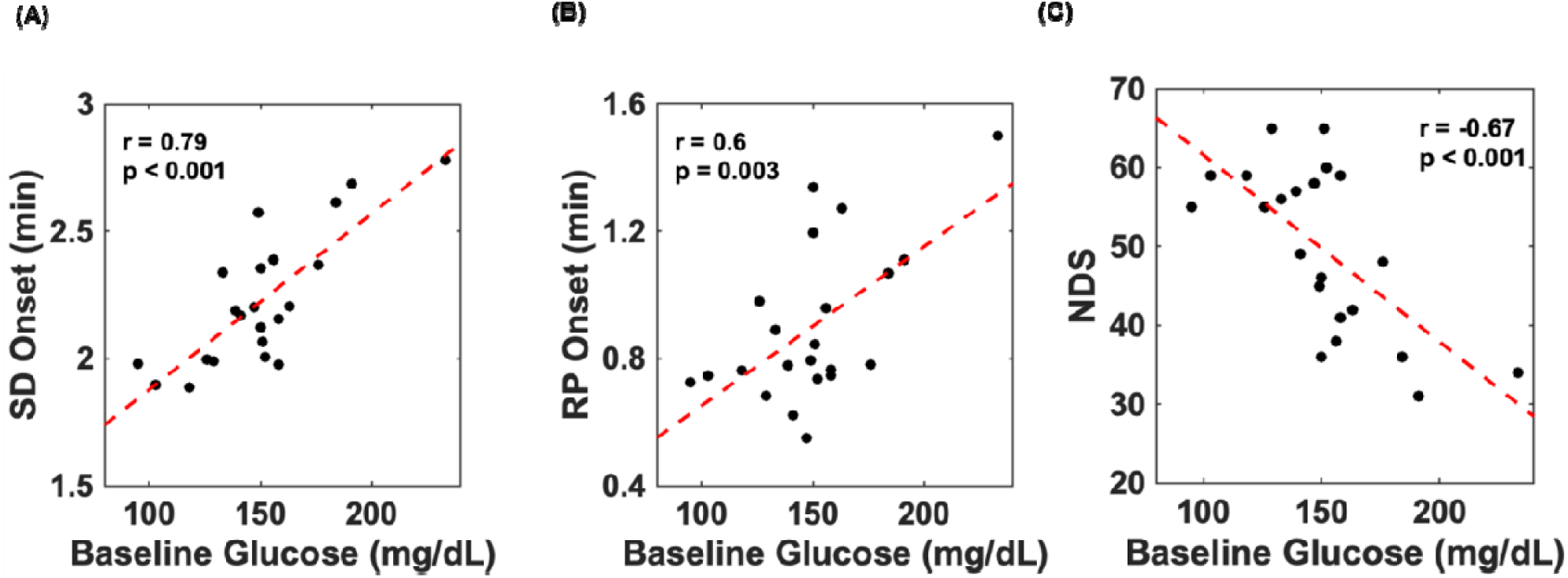
Baseline glucose correlates with SD onset, RP onset, and NDS.

